# Determining the Young’s Modulus of the Bacterial Cell Envelope

**DOI:** 10.1101/2024.04.04.588172

**Authors:** Junsung Lee, Karan Jha, Christine E. Harper, Wenyao Zhang, Malissa Ramsukh, Nikolaos Bouklas, Tobias Dörr, Peng Chen, Christopher J. Hernandez

## Abstract

Bacteria experience substantial physical forces in their natural environment including forces caused by osmotic pressure, growth in constrained spaces, and fluid shear. The cell envelope is the primary load-carrying structure of bacteria, but the mechanical properties of the cell envelope are poorly understood; reports of Young’s modulus of the cell envelope of *E. coli* are widely range from 2 MPa to 18 MPa. We have developed a microfluidic system to apply mechanical loads to hundreds of bacteria at once and demonstrated the utility of the approach for evaluating whole-cell stiffness. Here we extend this technique to determine Young’s modulus of the cell envelope of *E. coli* and of the pathogens *V. cholerae* and *S. aureus.* An optimization-based inverse finite element analysis was used to determine the cell envelope Young’s modulus from observed deformations. The Young’s modulus of the cell envelope was 2.06 ± 0.04 MPa for *E. coli*, 0.84 ± 0.02 MPa for *E. coli* treated with a chemical known to reduce cell stiffness, 0.12 ± 0.03 MPa for *V. cholerae*, and 1.52 ± 0.06 MPa for *S. aureus* (mean ± SD). The microfluidic approach allows examining hundreds of cells at once and is readily applied to Gram-negative and Gram-positive organisms as well as rod-shaped and cocci cells, allowing further examination of the structural causes of differences in cell envelope Young’s modulus among bacteria species and strains.

## 1. INTRODUCTION

Mechanical forces have a profound effect on cell physiology and survival. Most of our understanding in the field of cell biomechanics focuses on eukaryotes, primarily mammalian cells. However, other forms of life are more common: the total biomass of bacteria on Earth is 35 times greater than that of animals^1^. Bacteria experience substantial mechanical stimuli in the environment, including hydrostatic pressure, fluid shear stress, and adhesive forces^2–4^. In response to the physical environment, bacteria modify cell motility and/or initiate biofilm synthesis^5,6^. Furthermore, the mechanical properties of bacteria themselves may play a profound role in bacterial virulence: bacteria have been found to deform by as much as 80% to colonize sub micrometer-sized channels in bone^7^. We recently found that mechanical stress and strain within the cell envelope can influence the assembly and function of multicomponent efflux systems responsible for removing toxins^8^. As with other mechanosensitive mechanisms, the bacterial response to mechanical stimuli is likely mediated by the Young’s modulus (the material stiffness) of the bacterial cell envelope.

The small size of bacteria (∼1µm) makes it challenging to measure the mechanical properties of the cell envelope^4^. Several methods of applying mechanical loads to individual, live bacteria have been used to date. Atomic force microscopy involves the placement of probes in direct contact with the cell surface and provides force and deflection information that can be used to measure the Young’s modulus of whole cell (composite of the cell envelope and cytoplasm) or, with appropriate modeling, the Young’s modulus of cell envelope alone ^9–15^.

Alternatively, whole bacteria may be submitted to bending tests by securing one end of the cell, inducing filamentous growth, and applying a bending load using either optical tweezers or transverse fluid flow in a microfluidic chamber to determine flexural rigidity and calculate cell envelope Young’s modulus^16–19^. Lastly, the Young’s modulus of the cell envelope can be inferred by growing bacteria within an agarose gel of known stiffness by measuring the rate of cell elongation^20^. The Young’s modulus of the cell envelope of *E. coli* determined using these approaches spans from 2 MPa to 18 MPa (Table 1). While useful, these approaches have one or more of the following limitations: 1) they require the bacteria to be constrained to a surface (AFM, optical tweezers, gel encapsulation); 2) they require filamentous growth which alters bacterial physiology (bending experiments); and/or 3) the boundary conditions are poorly defined, for example, the underlying support for a cell submitted to an AFM probe^6^.

**Table 1.** Reports of the Young’s modulus of the cell envelope in *E. coli* are shown (mean ± SD). Measurement was performed on wet cells.

| Author | Method | Young's Modulus (MPa) | Species |
| --- | --- | --- | --- |
| Wang et al.<sup>19</sup> | Optical bending | $3.6 \pm 1.1^*$ | *E. coli* |
| Amir et al.<sup>16</sup> | Fluid bending | $6.4 \pm 1.9^*$ | *E. coli* |
| Auer et al.<sup>17</sup> | Fluid bending | $11.0 \pm 3.4^*$ | *E. coli* |
| Caspi<sup>18</sup> | Fluid bending | $4.4 \pm 1.3^*$ | *E. coli* |
| Deng et al.<sup>10</sup> | AFM indentation | $3.1 \pm 1.5^\ddagger$ | *E. coli* |
| Tuson et al.<sup>20</sup> | Gel encapsulation | $2.4 - 17.8^\ddagger$ | *E. coli* |

\*Young's modulus value (E) is calculated from reported flexural rigidity (EI) assuming $I = \pi r^3 t$
where $t = 33.70 \pm 2.90$ nm, $r = 0.42 \pm 0.04$ μm
‡Young's modulus value is calculated from deformation using cell envelope thickness, $t = 33.70 \pm 2.90$ nm

To explore the biomechanics and mechanobiology of bacteria without the limitations of the methods discussed above, our group developed a microfluidic approach we call “extrusion loading”^21^. Unlike other approaches, extrusion loading does not require permanent adhesion of bacteria to a surface and does not require the induction of filamentous growth. Furthermore, the extrusion loading system can examine hundreds of cells at once and is therefore much less labor-intensive than atomic force microscopy. During an extrusion loading experiment, bacteria are forced into submicron-scale tapered channels using fluid pressure. Extrusion loading can detect differences in whole cell stiffness: under the same magnitude of fluid pressure, less stiff cells travel further into a tapered channel than more stiff cells^21^. Although these prior studies have demonstrated the use of extrusion loading to detect differences in whole cell stiffness, and to study bacterial response to loading, the mechanical properties of the cell envelope were not determined. The Young’s modulus of the cell envelope is a major contributor to whole cell stiffness and likely influences mechanotransduction within the cell membrane or periplasm.

The long-term goal of this line of investigation is to determine the role of the mechanical properties of the bacterial cell envelope on cell physiology and survival under adverse conditions. Here we use optimization-based inverse finite element simulations of extrusion loading experiments to determine Young’s modulus of the cell envelope of *E. coli*, and the pathogens *V. cholerae* and *S. aureus*.

## 2. MATERIALS AND METHODS

### 2.1. Experimental Treatment: Extrusion Loading with Microfluidic Device

The extrusion loading approach was introduced in detail in prior work^21^ and is only briefly reviewed here. Extrusion loading involves forcing bacteria into submicron-scale tapered channels under fluid pressure. Extrusion loading is performed using microfluidic devices manufactured from fused silica using deep UV lithography^21^. A functioning extrusion loading device consists of 4-12 sets of tapered channels (5 channels per set) and a bypass channel (Figure 1a). Fluid flow through the bypass channel generates a pressure difference between the entry and exit of the tapered channels. Cells forced into the tapered chamber are deformed by the rigid channel walls (Figure 1b). Cells experiencing greater differential pressure travel further into the tapered channel and are more greatly deformed (Figure 1a). The distance traveled by the cells within the tapered channels and the deformed cell width are measured from bright field images (100x magnification).

**Figure 1.**
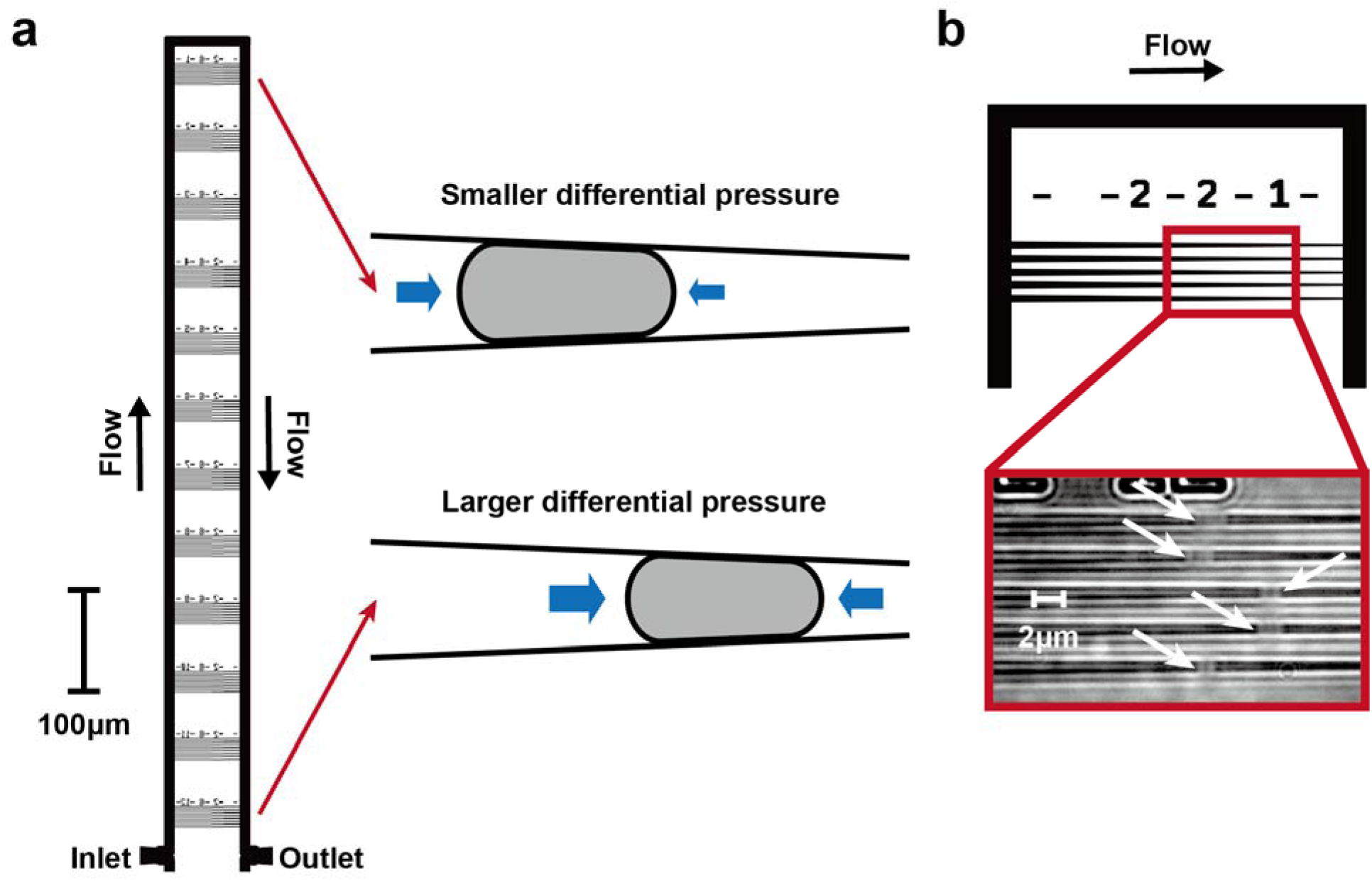
(a) An extrusion loading device is shown (10-16 devices are used in parallel in a single experiment). Bacteria in liquid media flow in through the inlet and either continue through the bypass channel or are trapped within one of the tapered channels. 12 sets of tapered channels are shown (5 channels/set), each set providing a distinct magnitude of differential pressure. A cell submitted to a greater magnitude of differential pressure undergoes more deformation (reduction in cell width and further distance travelled into the tapered channel). (b) A bright field image of bacteria (depicted with white arrows) trapped inside of the tapered channels is shown.

An extrusion loading experiment was performed as follows. The microfluidic device was prewet with liquid media. Cells were grown to exponential phase and flowed into the microfluidic device at room temperature using a fluid pressure of 60 kPa using a fluid pump system (PneuWave Pump, CorSolutions, Ithaca, NY, USA). In a single experiment, as many as 720 cells were submitted to extrusion loading at differential pressure values ranging from 0.3 kPa to 13 kPa. The differential pressure values were determined using hydraulic circuit calculations and were specific to each device/experiment (device dimensions and filling pattern of the tapered channels were taken into account in the hydraulic circuit calculations)^8,21,22^.

We analyzed data from four experiments (Table 2): three examining *E. coli* and *S. aureus* were performed for this study and one involving *V. cholerae* from a previously published study^22^. In the first series of experiments deformation of *E. coli* under extrusion loading was observed. In the second series of experiments *E. coli* were treated with the antimicrobial molecule A22 (10 µg/mL) for 20 minutes prior to extrusion loading. A22 depolymerizes the MreB protein within *E. coli*^23^. MreB is a “shape determining” protein crucial for maintenance of rod shape^24^. We previously found that depolymerization of the MreB protein led to reduced whole cell stiffness^21^. Hence, A22 treated *E. coli* submitted to extrusion loading are expected to display a reduced cell envelope Young’s modulus. The third series of experiments were conducted on rod-shaped *V. cholerae* created by deleting the periplasmic protein *crvA*, which is necessary for inducing curvature in *V. cholerae*^25^. In the fourth series of experiments, *S. aureus* was submitted to extrusion loading. Experiments using *E. coli* and *V. cholerae* also involved single molecule tracking using fluorescence and were performed with cells suspended in M9 minimal medium to avoid autofluorescence in the standard growth media, LB. Analysis of *S. aureus* was performed in nutrient rich media (TSB) that did not interfere with our analyses.

**Table 2.**
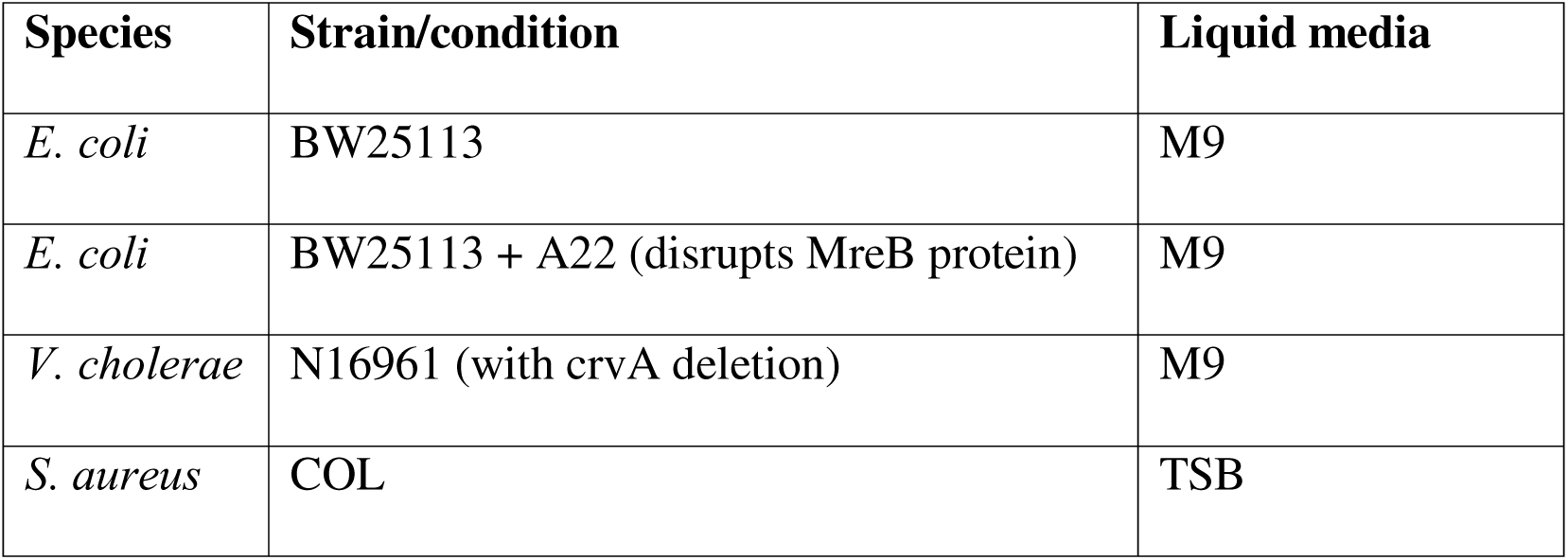
Study groups used to evaluate cell envelope Young’s modulus are shown.

The relationship between applied fluid pressure and distance traveled into the tapered channels during extrusion loading is nonlinear^21^. For measurements of Young’s modulus, we used cells loaded within the linear range, which did not include cells with minimal deformation (deformed cell width in tapered channels similar to free-floating cell width) or cells too close to the taper exit (thereby preventing cells that slip out of the tapered channels from influencing mean and standard deviation in measurement of deformed cell width in tapered channels).

Finite element model of bacterial cell envelope under extrusion loading Here we used experimental data from the four series of experiments each consisting of at least 3 replicates, leading to examination of 258 – 536 cells. We used an optimization-based inverse finite element analysis to determine the Young’s modulus of the bacterial cell envelope from the experimental findings. In an inverse finite element model the unknown material properties and loading conditions (in this case cell envelope Young’s modulus and cell internal pressure when the cell is submitted to extrusion loading) were assigned initial values, a finite element simulation was performed, and the resulting deformations were compared to whole cell deformation at different differential pressure magnitudes. The values of unknown parameters were then adjusted using an optimization algorithm and the finite element analysis was repeated. The process continued until the sum of squared error between the finite element model and the experimental results is minimized. The simulations were repeated with different initial parameters to ensure that the final optimized result is not a local minimum.

Finite element models of bacteria under extrusion loading were generated using ABAQUS (version 2019, Dassault Systèmes, Vélizy-Villacoublay, France). The cell envelope was modeled as a homogenous material (i.e., the cell wall, periplasm, outer membrane and inner membrane are considered a single composite material). Rod-shaped bacteria (*E. coli* and *V. cholerae*) were modeled as a pressure vessel with a cylindrical trunk and hemispherical caps at two ends (Figure 2a and 2b) and spherical-shaped bacteria (*S. aureus*) were modeled as a spherical pressure vessel (Figure 2c). The model used linear elastic solid axisymmetric elements, consistent with prior work modeling the mechanical properties of the bacterial cell envelope^26^.

**Figure 2.**
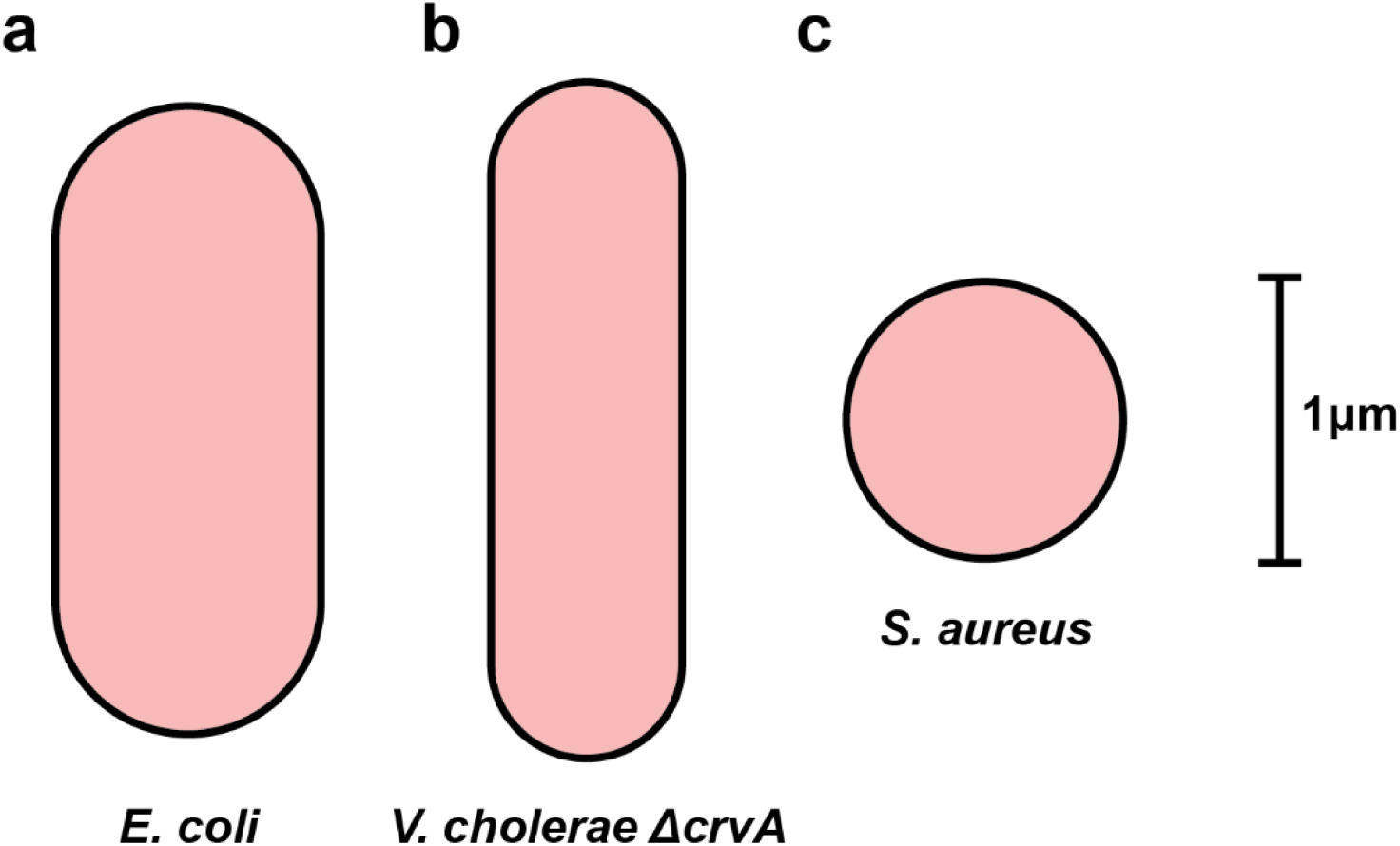
The average size and shape for (a) *E. coli* (b) *V. cholerae* Δ*crvA* (to achieve a rod-like shape), and (c) *S. aureus* is shown. Each cell was modeled as a pressure vessel.

The models were composed of four-node axisymmetric bilinear solid elements (CAX4) with linear elasticity and geometric nonlinearity (Figure 3a). Mesh convergence test showed that increasing the number of elements has a negligible effect in the simulation results (Figure S1).

**Figure 3.**
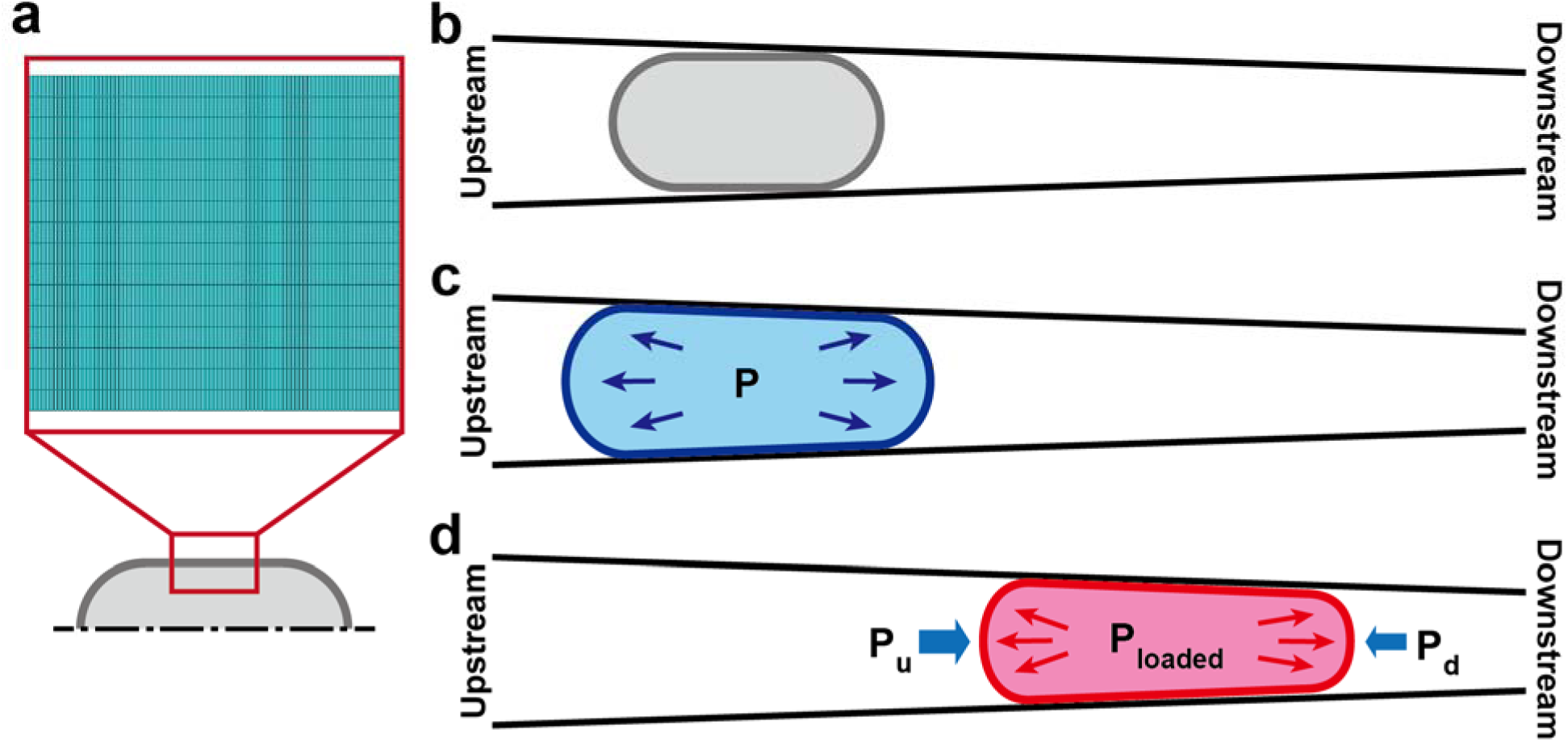
(a) The cell envelope was modeled axisymmetric (inset: finite element mesh). The finite element simulation involved three steps: (b) the hypothetical unstressed state of the cell envelope without turgor; (c) internal pressure, P, is applied to create contact with the channel walls; and (d) application of fluidic pressure caused by extrusion loading (P_U_ and P_d_) along with an increase in internal pressure to P_lOaded_.

The cell envelope of rod-shaped bacteria displays transverse isotropy in the trunk section, with Young’s modulus in the hoop direction twice as large as that in the axial direction (anisotropy ratio of 2), consistent with the orientation of peptidoglycan strands and the orientation of the MreB protein^10,27^. Preliminary simulations demonstrated that variation in the value of anisotropy ratio has negligible effects on the finite element simulation results (Figure S2). The Young’s modulus in the cap sections of rod-shaped bacteria was set as the average Young’s modulus of the trunk section since the peptidoglycan strands in the cap in this region are aligned randomly^28^. *S. aureus* was modeled as a spherical pressure vessel with isotropic cell envelope. A Poisson’s ratio of 0.3 was used, as it is the mean value determined by molecular dynamics simulations (which ranged from 0.22 to 0.67) in prior experimental studies^10,20,27,29^.

Preliminary simulations demonstrated that variation in the value of Poisson’s ratio has a negligible effect on the results obtained with the model (Figure S2). The Young’s moduli in the axial direction (Ea) for rod-shaped bacteria and Young’s modulus (E) for spherical shaped bacteria were unknown and used as a free variable in the inverse finite element model.

The geometry of the cell in its turgid and unstressed (no turgor) state was determined from studies examining cell width and the percent decrease in size caused by rapid loss of turgor and plasmolysis in *E. coli*, *V. cholerae*^30^ and *S. aureus*^31^; cell width shows little variation within a population of bacteria^32^ (see Table 3 for the values of geometry of the unstressed cell and see Supplementary Method and Figure S3 in the Supporting Information for the detailed calculation of the geometry of unstressed cell). In this study, we relied on existing literature that provided numerical values for the thickness of the cell envelope in each species^31,33^ (Table 3). Even though the information regarding cell envelope thickness was not available for each bacterial strain examined, preliminary simulations demonstrated that variation in the value of cell envelope thickness within a reasonable range has small effect on the finite element simulation results (Figure S2).

**Table 3.**
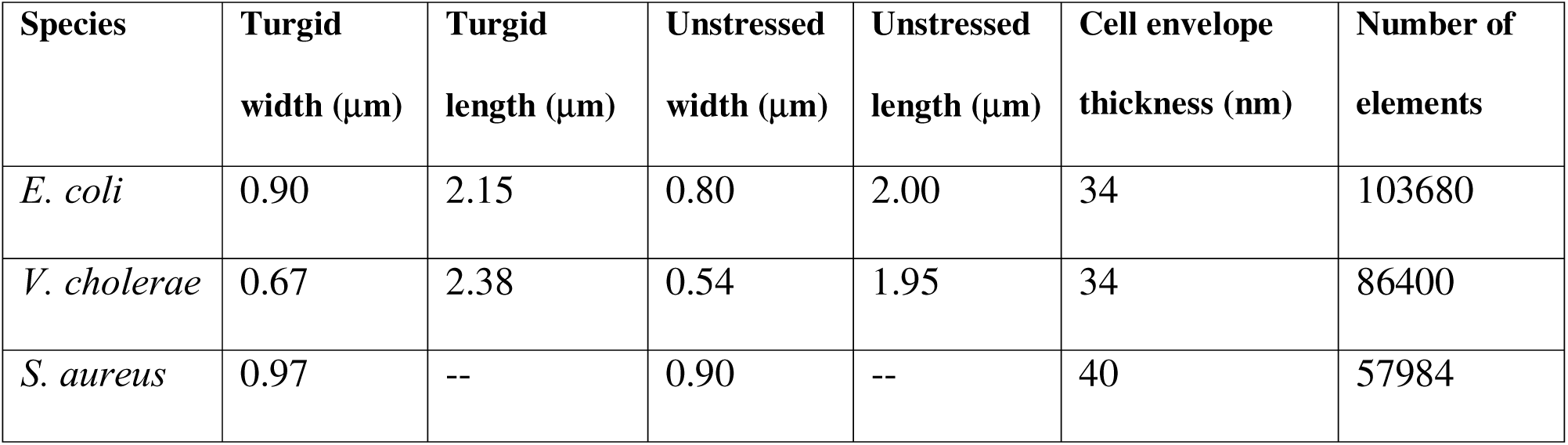
Characteristics of the finite element models of each species are shown.

We simulated extrusion loading using the standard finite element modeling approach for an elastic pressure vessel submitted to external loads^34^: Simulations began with the cell envelope in an unstressed state followed by the application of turgor pressure, and then followed by external loads from extrusion loading. This two-step process makes it possible to determine the absolute mechanical stress states of the cell envelope due to the extrusion loading, not only the difference in mechanical stress from that caused by turgor alone. The unstressed cell envelope (in the absence of turgor pressure) was placed within the tapered channel walls so that the cell envelope was in contact with the rigid wall at a single point (Figure 3b). Friction at contact between the cell and the channel wall was neglected, as is done when modeling micropipette aspiration^35–37^. The cell envelope was then inflated to bring the cell into contact with the channel walls by applying an internal pressure (P). The simulation results were insensitive to this initial value of internal pressure as long as the trunk section of the inflated cell envelope is fully in contact with the channel walls (see Supplementary Discussion and Figure S4 in the Supporting Information for more details). Additionally, a pin boundary condition was used to prevent rigid body motion of the cell during the inflation (Figure 3c).

Once the inflated cell was in contact with the walls, the pin boundary condition was removed, a differential pressure (P_U_ and P_d_) was applied outside of the cell envelope, and the internal cell pressure was increased to a new value, P_lOaded_ (Figure 3d). Analytical models of cells submitted to extrusion loading demonstrated that loading is associated with increased internal pressure, most likely due to increases in cell osmolarity resulting from a reduction in cell volume caused by water loss^8^. However, the magnitude of the increase in turgor pressure is unknown. The internal cell pressure when loaded (P_lOaded_) was therefore an unknown variable and solved for using the inverse finite element model. The differential pressure values (P_U_ and P_d_) were specific to each of the different pressure levels within the device determined using hydraulic circuit calculations^8,21^. The application of the differential pressure caused stepwise rigid body motion of the cell toward the downstream end of the tapered channel until a final deformed cell width was reached.

A Python script (version 3.9.7, Python Software Foundation, Fredericksburg, VA, USA) was used to generate input files for ABAQUS for each iteration of the optimization process. Each finite element simulation took 20-60 minutes using a 3.00 GHz processor with 12 cores and 6 parallel threads and 64 gigabytes of RAM. Each iteration of the inverse finite element analysis (considering one pair of values for E_a_ and P_lOaded_) consisted of 4–11 nonlinear axisymmetric simulations, one for each magnitude of differential pressure applied in the simulated experiment.

### 2.2. Determination of cell envelope Young’s modulus using an optimization-based inverse finite element analysis

The unknown variables in the inverse model included the Young’s modulus (E_a_, Young’s modulus in the axial direction, for rod-shaped cells and E for spherical cells) and internal cell pressure when loaded (P_lOaded_). A least squares optimization algorithm was used to adjust the two unknowns between optimization iterations. The combination of cell envelope Young’s modulus (E_a_ or E) and internal cell pressure when loaded (P_lOaded_) that resulted in finite element simulations most consistent with the experimental results was determined by minimizing the least squared error objective function:

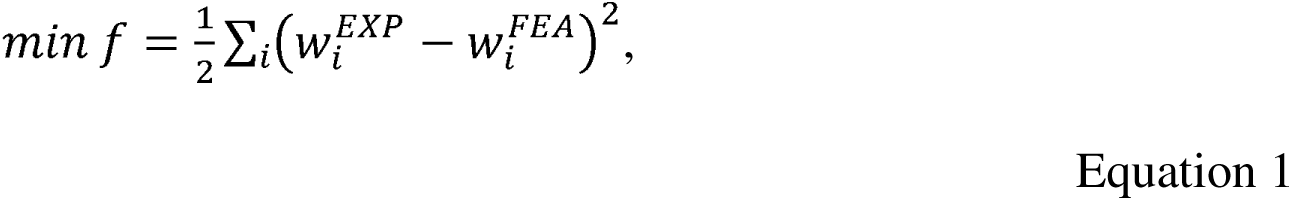

where *i* is the differential pressure level experienced by the cell (4-11 different differential pressure vessels per species/condition), w^EXP^ is the mean value of the deformed cell widths observed in the extrusion loading experiments at pressure level i; and w^FEA^ is the deformed cell width from the finite element simulation for pressure level i. To solve the objective function, a nonlinear least squares optimization function using the Trust Region Reflective algorithm was implemented in Python using scipy.optimize package (version 1.7.3)^38,39^.

The optimization process started with a pair of initial guesses for the free variables: Young’s modulus () and internal cell pressure when loaded (). A Python script generated ABAQUS input files for the observed differential pressures in the experimental data and executed finite element simulations. After the simulations were completed, the script determined the deformed cell width values from each simulation and determined error between the simulated width and the mean deformed cell width from the experiments at each differential pressure level. The optimization algorithm determines if the sum of squared error is minimized. If the sum of squared error is not minimized the optimization algorithm calculates a new pair of free variables (E_a_, P_lOaded_) for rod-shaped cells and (E, P_lOaded_) for spherical cells, and this process was repeated until the optimization algorithm has successfully minimized the sum of squared error (Figure 4). To ensure that the final values identified by the optimization are not a local minimum, the optimization was performed with different pairs of initial guesses for Young’s modulus (E_a_ar E) and the internal cell pressure when loaded (P_lOaded_) spanning the range of expected values (Table 4).

**Table 4.** The values of initial guesses used in the optimization are shown. Three different initial guesses were used to confirm the global minimum.

| Species | Initial guess pair ($E_a, P_{loaded}$) | | |
| --- | --- | --- | --- |
| *E. coli* | (0.1 MPa, 0.03 MPa) | (4 MPa, 0.2 MPa) | (20 MPa, 0.5 MPa) |
| *V. cholerae* | (0.1 MPa, 0.03 MPa) | (4 MPa, 0.3 MPa) | (12 MPa, 0.9 MPa) |
| *S. aureus* | (0.4 MPa, 0.06 MPa) | (4 MPa, 0.2 MPa) | (20 MPa, 0.5 MPa) |

**Figure 4.**
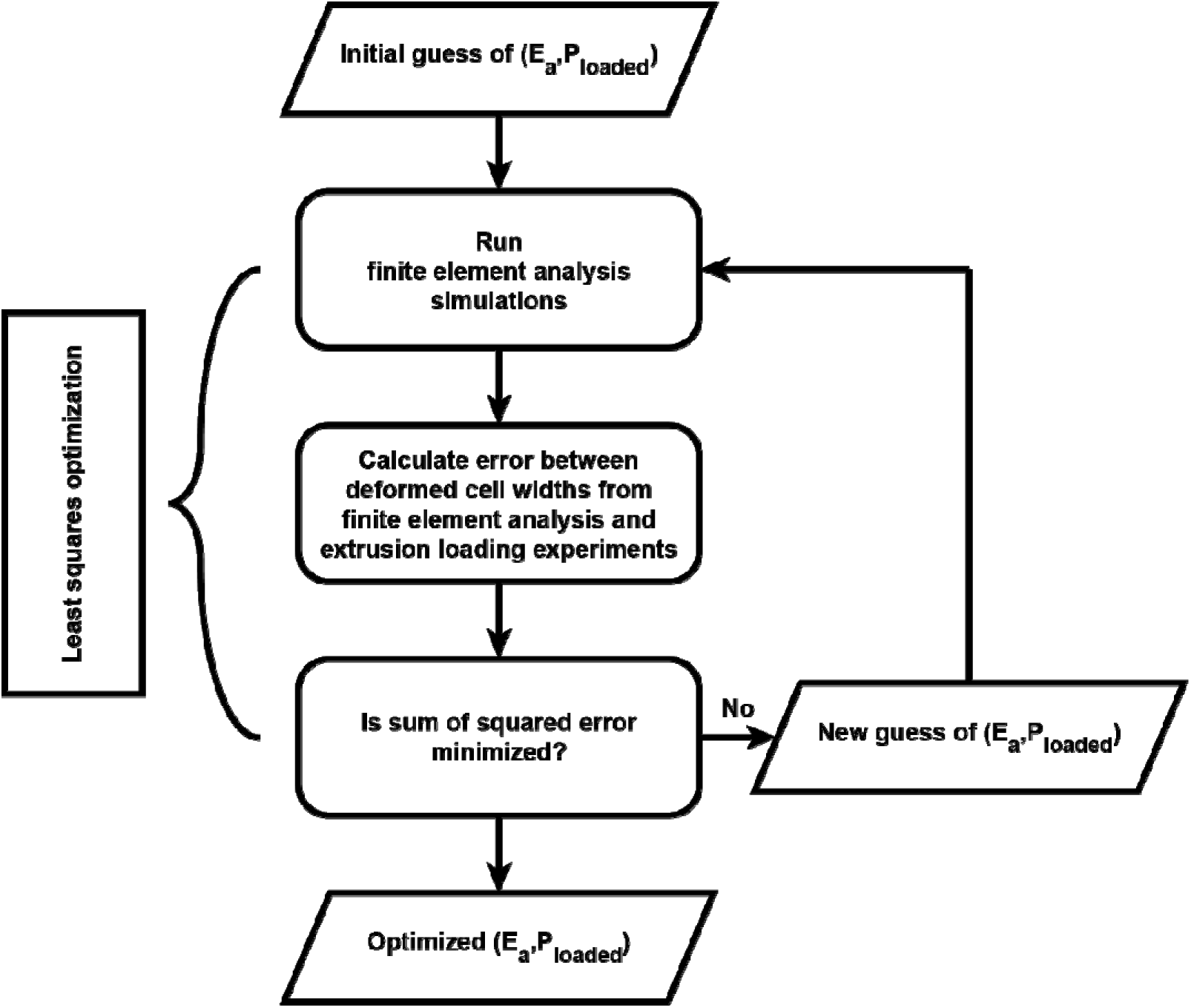
A flow chart of the optimization-based inverse finite element modeling method is shown.

Using the Young’s modulus is obtained using the inverse finite element analysis, we were also able to estimate the turgor pressure of the cell in liquid media by creating a mechanical model using the turgid cell width (measured directly here) and the percent reduction in cell dimensions reported when turgor pressure is removed through plasmolysis for each of the three bacterial species modeled (see^30,31^ and Supplementary Method in the Supporting Information for more details).

## 3. RESULTS

Cells deformed more under greater differential pressure. *E. coli* exhibited the largest cell width of all tested bacteria under all magnitudes of differential pressure and depolymerization of MreB protein (by A22 treatment) increased the deformation of *E. coli* by 20%. *V. cholerae* showed the smallest deformed cell width among the four groups. *S. aureus* showed smaller deformed cell widths than *E. coli* treated with A22 but larger deformed cell widths than *V. cholerae* (Figure 5).

**Figure 5.**
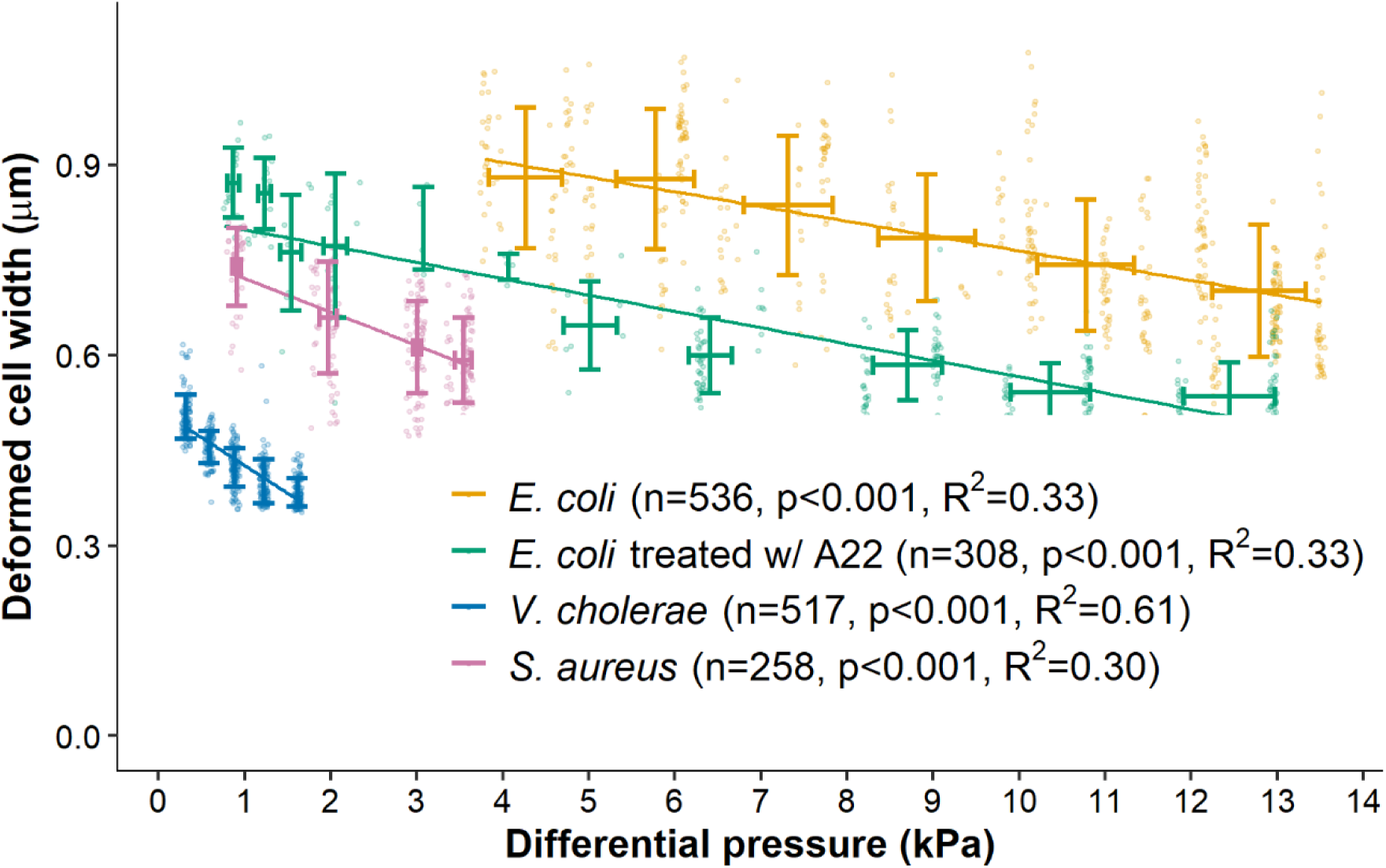
The deformed cell width and differential pressure of the experimental data sets used in this analysis are shown. Data are binned into a finite number of differential pressure values to facilitate the inverse finite element (data points and mean ± standard deviation in both axes).

Parentheses contain the number of datapoints and p-value and R^2^ value of the linear regression. The range of differential pressure differs among the experimental groups after accounting for variations in microfluidic device dimensions and patterns of channel occupancy observed in each experiment.

Each inverse finite element model analysis involved 5-10 iterations totaling 120-330 finite element simulations. For each species/condition, optimizations with the three initial guesses of Young’s modulus (E_a_ar E) converged to a single value, demonstrating that the optimized values were unlikely to be the local minima (Figure 6a-d). The Young’s modulus of the cell envelope of *E. coli* determined in the model was 2.06 ± 0.04 MPa (mean ± SD of optimization results with three different initial guesses) (Figure 6a). The Young’s modulus of the cell envelope of A22 treated *E. coli* was 0.84 ± 0.02 MPa (Figure 6b); 59.2% smaller than that in untreated *E. coli*. The Young’s modulus of the cell envelope of *V. cholerae* was 0.12 ± 0.03 MPa (Figure 6c). The Young’s modulus of the cell envelope of *S. aureus* was 1.52 ± 0.06 MPa (Figure 6d). Finite element simulations using the above cell envelope Young’s modulus values resulted in the deformed cell width with an average error of 1.3% compared to the experimental observations (Figure 6e, 6f).

**Figure 6.**
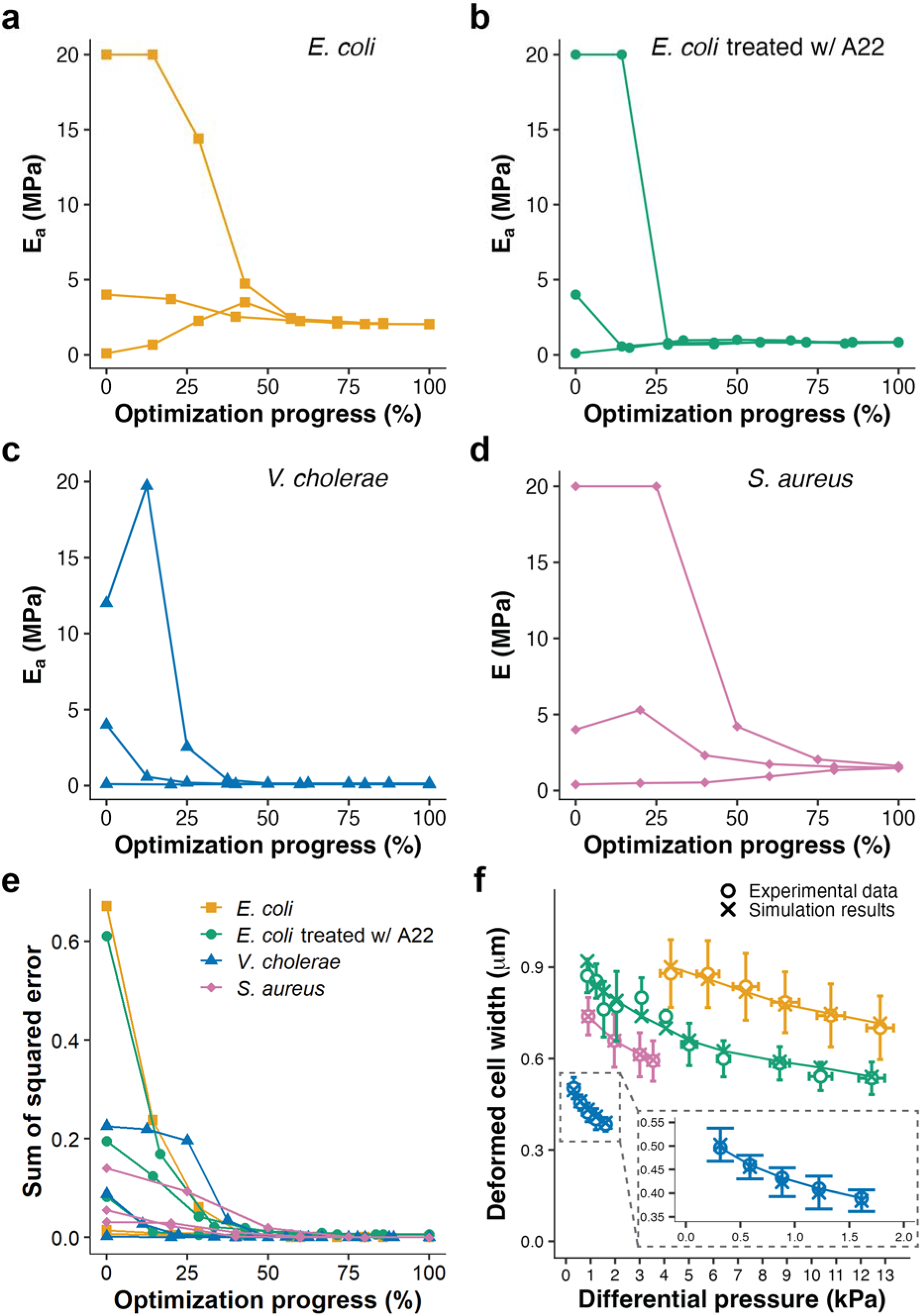
The convergence histories of optimization from different initial guesses are shown for the Young’s moduli for (a) *E. coli*, (b) *E. coli* treated with A22, (c) *S. aureus,* and (d) *V. cholerae.* Each data point represents one iteration (e) Sum of squared error minimized with the progress of the optimization is shown. (f) Comparison of the experimental data to the results of the finite element simulations is shown (the color scheme of the current plot remains consistent with that of the preceding plot and the inset provides a magnified representation of the region demarcated in the main plot).

The width of a turgid cell shows relatively little variation among cells in the same media^32^. As a result, there is a relationship between the Young’s modulus and the turgor pressure of the cell which we determined using thin-walled pressure vessel calculations (see Supplementary Method in the Supporting Information for more details). The cell envelope Young’s modulus values determined above resulted in the estimates of turgor pressure of 45.4 ± 0.8 kPa for *E. coli*, 9.4 ± 2.0 kPa for *V. cholerae,* and 27.4 ± 1.1 kPa for *S. aureus.* Each of these values are significantly different from one another (p<0.01, Bonferroni multiple comparisons).

The stress states within the cell envelope under extrusion loading indicate that greater differential pressure is associated with greater octahedral shear stress within the cell envelope, while the axial stresses (along the length of the rod-like cell), hoop stresses (circumferential in rod-like cells), and radial stresses (perpendicular to the cell envelope surface) are reduced at greater differential pressure (Figure 7). Shear stresses did not show significant change with increasing differential pressure.

**Figure 7.**
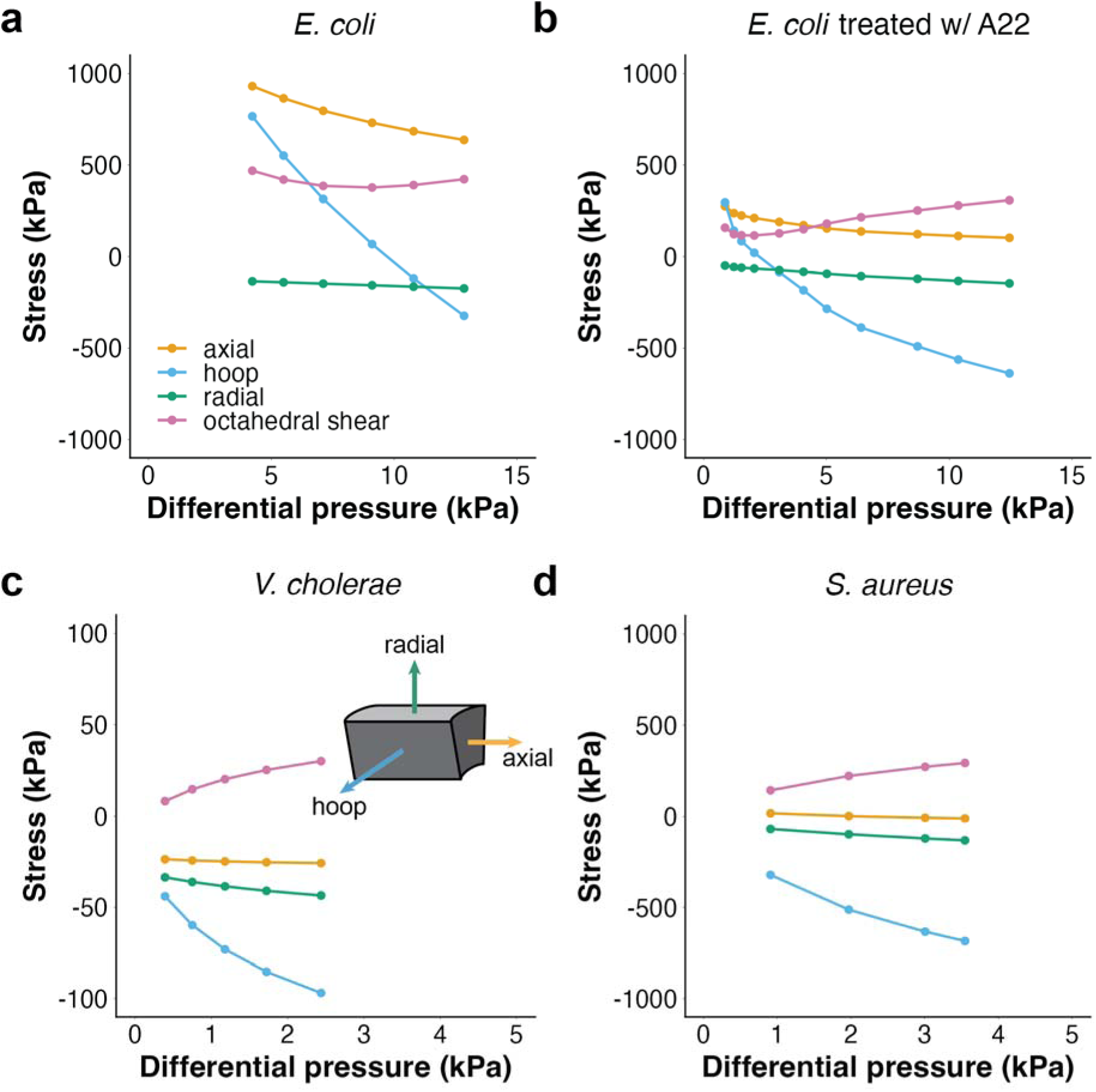
Mechanical stress in the cell envelope under extrusion loading determined in the finite element model is shown for (a) *E. coli,* (b) *E. coli* treated with A22, (c) *S. aureus*, and (d) *V. cholerae* (the inset shows the direction of the stress on an element in the cylindrical coordinate).

## 4. DISCUSSION

Here we demonstrated a method of determining the Young’s modulus of the bacterial cell envelope from experiments using extrusion loading applied using microfluidic devices. We applied the approach to estimate the Young’s moduli of the cell envelopes in four different species/conditions. Our findings demonstrate large variability in the cell envelope Young’s modulus among species.

The Young’s modulus of the cell envelope of *E. coli* determined in this study (mean value of 2.06 MPa) is slightly smaller than the previously reported range (2.4-17.8 MPa, Table 1). Our result is most comparable to previous AFM indentation experiments done by Deng et al. (reporting a value of 3.1 ± 1.5 MPa)^10^.

Inactivation of the structural protein MreB using the A22 antibiotic resulted in a reduction of cell envelope Young’s modulus by 59.2% (mean value of 0.84 MPa). The MreB protein in *E. coli* forms short circumferential strands on the inner membrane^40,41^. Inactivation of the MreB protein using A22, removes constraints on cell envelope growth, resulting in the growth of cells that are no longer rod-like. Hence, the MreB protein is referred to as a “shape determining” protein. Our finding that the Young’s modulus of the cell envelope in *E. coli* is reduced after depolymerization of the MreB protein is consistent with prior observations using bending^42^ and our prior observations using extrusion loading^21^. However, how MreB contributes to the mechanical performance of the cell envelope is not yet understood. MreB may contribute to the mechanical properties of *E. coli* by regulating the movement of Penicillin-binding protein 2 (PBP2) thereby establishing the orientation of peptidoglycan strands within the cell envelope during cell wall synthesis. Alternatively, the stiffness of the MreB protein itself may regulate the mechanical performance of the cell envelope in the same way that stiffening components are used on thin-walled pressure vessels.

The Young’s modulus of the cell envelope of *V. cholerae* measured in this study (mean value of 0.12 MPa) is more than an order of magnitude smaller than that seen for *E. coli* (mean value of 2.06 MPa). The considerably smaller Young’s modulus of *V. cholerae* was surprising but unlikely to be a result of experimental error or the small size of *V. cholerae* (different cell size was considered in the finite element models, see Table 3). While we are not aware of prior work measuring the Young’s modulus of the cell envelope of *V. cholerae*, plasmolysis induced by hyperosmotic shock caused much greater deformation of *V. cholerae* (21.8 ± 2.6%, mean ± SD) than *E. coli* (9.6 ± 2.9%)^30^. Such a difference in deformation after sudden removal of turgor would be consistent with a much smaller Young’s modulus (possible) or a much larger turgor pressure (which we believe to be unlikely, see below). Further investigation is required to understand why the Young’s modulus of the cell envelope of *V. cholerae* is so much smaller than that of *E. coli*. Some possible explanations include a lower peptidoglycan density (or degree of crosslinking) and/or differences in outer membrane properties. Peptidoglycan forms a cross-linked polymeric meshwork that stabilizes the cell and provides structural integrity^43^. In principle, a thinner cell envelope would result in a smaller Young’s modulus of the cell envelope. While the thickness of the cell envelope of *E. coli* and *S. aureus* have been reported^31,33^, we are not aware of such measurements made on *V. cholerae*. A further analysis on the relationship between cell envelope thickness and cell deformation (see Supplementary Discussion and Figure S5 in the Supporting Information for more details) suggests that the thickness of the cell envelope of *V. cholerae* would have to be 20 times less than that of *E. coli* for the simulations to achieve a cell envelope Young’s modulus similar to that of *E. coli*. As such a cell envelope thickness is smaller than the cell wall, the thickness of the cell envelope alone is unlikely to explain the disparity in the Young’s moduli of *E. coli* and *V. cholerae*. Alternatively, it is possible that there are differences in the constituents of the cell envelope in *V. cholerae,* such as a lower density of peptidoglycan or a less mechanically robust outer membrane. If confirmed in subsequent studies, the lower Young’s modulus of *V. cholerae* may provide survival advantages given the broader environmental habitats of *V. cholerae* (host gastrointestinal tract and freshwater rivers) and during treatment with cell wall inhibitors.

The Young’s modulus of *S. aureus* determined in this study (mean value of 1.52 MPa) is slightly smaller than that seen in *E. coli*, which may seem surprising since Gram-positive bacteria have a much thicker peptidoglycan cell wall than Gram-negative bacteria and is therefore widely believed to be stiffer^43,44^. However, our measurement of the Young’s modulus of the cell envelope of *S. aureus* is well within the range of prior reports from studies using atomic force microscopy (0.57-1.8 MPa) which are also smaller than reports of the Young’s modulus in *E. coli* determined using AFM (see Table 1). Hence, our findings provide further support for the idea that the Young’s modulus of *S. aureus* is slightly smaller than that of *E. coli*. One possible explanation for the smaller Young’s modulus of the cell envelope of *S. aureus* is that the outer membrane of *E. coli* is a major contributor to the mechanical function of the cell envelope^30^, potentially making up for the much thinner peptidoglycan cell wall in *E. coli*.

There are several strengths to our methods that lend confidence in our results. The inverse finite element model converged on the same Young’s modulus value in each of the conditions modeled, giving confidence in the values determined by the optimization-based inverse finite element analysis. We attribute some of the differences between our observations and prior work to limitations in experimental approaches and computational analyses that our extrusion loading method solves. For example, studies using atomic force microscopy to determine cell envelope Young’s modulus required bacteria to be constrained on a surface. This experimental condition makes it difficult to define boundary conditions at the contact between the cell and the underlying substate, potentially generating errors when using whole cell deformations to determine the cell envelope Young’s modulus. Furthermore, studies using AFM each use different techniques to attach the cells on a surface^9,10,13,14^, potentially leading to variability in the contact between the cell and its substrate. Fluid bending experiments require filamentous growth of bacteria which may change cell physiology and potentially influence measurements or limit the types of bacteria that can be evaluated with the fluid bending approach. A major limitation of the gel encapsulation method is that applied forces are generated by cell elongation. Hence, the Young’s modulus determined from gel encapsulation is influenced by rates of cell elongation, which are slowed by the presence of the gel.

Our approach also provides estimates for the turgor pressure of the cell. Turgor pressure is the primary source of mechanical stress in bacteria and greatly influences cell growth and division^45^. Measurement of turgor pressure of bacteria has been hindered by the small size of bacteria^45,46^. In our work, turgor pressure is estimated using the Young’s modulus determined from the inverse finite element model, the reduction in cell envelope geometry reported following plasmolysis^30,31^ and the observed turgid cell width. The turgor pressure of *E. coli* estimated from our work (mean value of 45.4 kPa) is similar to that determined from recent AFM experiments (30 kPa^10^). In contrast, our estimate of turgor pressure in *S. aureus* (mean value of 27.4 kPa) is smaller than that seen in *E. coli* and substantially smaller than common estimates of the turgor pressure of *S. aureus* (typically between 2-3 MPa^31^). However, others have reported that the deformations of *S. aureus* following removal of turgor pressure during hyperosmotic shock (6.7 ± 1.1%, mean ± SD^31,47^) is smaller than that seen with *E. coli* (9.6 ± 2.9%^45^). If the Young’s modulus of the cell envelope of *S. aureus* is also smaller than that of *E. coli* (as reported by others and also found in this study), a smaller turgor pressure of *S. aureus* might be expected. Similarly, the turgor pressure estimated for *V. cholerae* (mean value of 9.3 kPa) is substantially smaller than that seen for *E. coli*. Previous work has shown that *V.* cholerae responds to removal of its cell wall (e.g. after exposure to beta-lactam antibiotics) by forming cell wall-deficient spheroplasts, while *E. coli* lyses rapidly under the same conditions^48^. Reduced turgor could in principle be the underlying cause of *V. cholerae*’s ability to retain structural integrity in the absence of a cell wall.

There are some limitations that must be considered when interpreting our results. First, we have modeled the cell envelope as a homogenous material, thereby lumping together the inner membrane, periplasm, cell wall and outer membrane as a continuum. This assumption is the current standard in mechanical modeling of bacterial cell envelopes but does not address the underlying physics of each constituent. Additionally, we have modeled the cell envelope as a linear elastic solid, although the cell envelope is a soft material and therefore likely to display material nonlinearities (such as viscoelasticity).However, the linear elastic solid assumption is the current standard for modeling the bacterial cell envelope^26^ and there are no existing experimental data to suggest an appropriate nonlinear model. Additionally, our simulations were simplified to use a single value for the internal cell pressure (P_lOaded_) under loading at all magnitudes of differential pressure (see Figure S6 for optimized P_lOaded_ values). Our previous analyses indicate that P_lOaded_ may be related to the decrease in cell volume during extrusion loading which is more pronounced at larger magnitudes of extrusion loading and therefore causes increases in axial stress^8^. Upon closer analysis, however, the changes in cell volume during extrusion loading are not well understood because current experiments do not evaluate cell deformation in the corners of the rectangular channels within the microfluidic device.

Without a better understanding of the changes in cell volume during extrusion loading it is difficult to establish the appropriate changes in P_lOaded_ during extrusion loading. It is possible that slight improvements in the estimates of Young’s modulus could be achieved if changes in P_lOaded_ at each applied differential pressure were better understood or integrated as an unknown into the optimization algorithm, which we did not consider here because it would increase the number of finite element simulations by 4-11 times (i.e. a different value of P_lOaded_ for each differential pressure magnitude). Since there is additional deformation of the cells into the corners of the rectangular microfluidic device the estimation of Young’s modulus in our study, which based on the assumption of a conical channel shape, represents a conservative estimate of the true value. However, it is important to acknowledge that any error in Young’s modulus estimation is mitigated by assuming frictionless contact. Although the conical channel restricts cell travel, the frictionless assumption enables the cells to travel further in the finite element simulations. Thus, these compensatory effects reduce the error in Young’s modulus estimation. To understand and address these limitations and thereby improve evaluation of Young’s modulus, future studies should incorporate the 3D shape of loaded cells within microfluidic devices and employ comprehensive 3D modeling approaches.

## 5. CONCLUSIONS

The extrusion loading approach is advantageous in that it provides a means of applying mechanical loads to hundreds of bacteria without requiring attachment to a surface or changing physiology to cause filamentous growth. Hence our methodology provides a means to quantitatively study the influences of cell envelope mechanical properties on cell physiology and survival following exposure to antibiotics and other toxins. Also, this study demonstrated the quantitative difference in mechanical stiffness of cell envelope among different species of bacteria including Gram-negative and Gram-positive organisms. While the existing knowledge concerning the mechanical effects of structural distinctions among various species of bacteria remains limited, this study facilitates an investigation into the disparities in mechanical stiffness within the cell envelope, which may arise from physiological and structural variations among different species. Quantitative assessment of the mechanical properties of the cell envelope makes it possible to interrogate changes more thoroughly in the cell envelope caused by antibiotics and other factors that influence bacterial survival.

## SUPPORTING INFORMATION AVAILABLE

The following file is available free of charge.

SI_E_of_bacterial_cell_envelope.pdf: mesh convergence of the finite element model; sensitivity analyses of anisotropy ratio, Poisson’s ratio, and cell envelope thickness; detailed calculations for geometry of cells in unstressed state; detailed derivation of the relationship between Young’s modulus of cell envelope and turgor pressure; histograms of measured cell dimensions; sensitivity analysis on the internal cell pressure; additional discussion on the effect of cell envelope thickness on cell envelope Young’s modulus determined using the finite element model; and convergence histories of optimization from different initial guesses of the internal cell pressure under extrusion loading.

## TABLE OF CONTENTS GRAPHIC

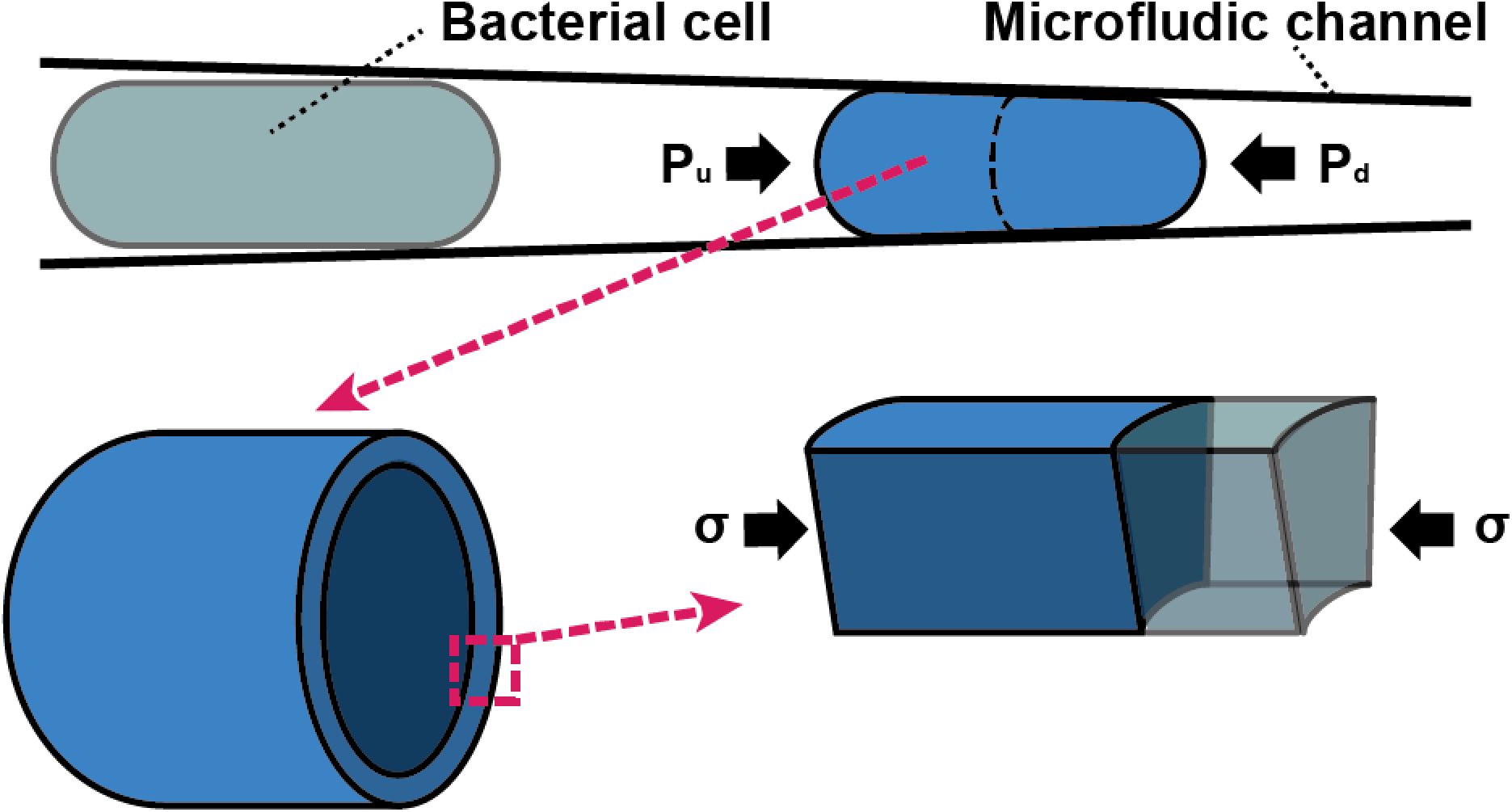

